# The Taxonomic Status and Phylogenetic Relationships of *Toxorhynchites* (Diptera: Culicidae) Species from Panama

**DOI:** 10.64898/2026.08.10.744079

**Authors:** Bennett-Vaz Richard, Mabelle Chong, Ambar L. Rojas, Yosiat Vega, Jose R. Loaiza

**Affiliations:** División de Una Sola Salud, Instituto de Investigaciones Científicas y Servicios de Alta Tecnología (INDICASAT AIP), Panamá, República de Panamá; Programa Centroamericano de Maestría en Entomología, Universidad de Panamá, Campus Octavio Méndez Pereira, Avenida Transístmica, República de Panamá; Smithsonian Tropical Research Institute, Apartado 0843-03092, Panama City, Panama

**Keywords:** Molecular barcoding, phylogenetics, predatory larvae, non-biting females, species delimitation

## Abstract

Mosquitoes in the Toxorhynchites genus (Theobald, 1901) are specialized predators of container-breeding mosquito larvae, including major Aedes disease vectors. Despite their biological control potential, species boundaries and phylogenetic relationships remain poorly understood. Utilizing country-wide sampling across Panama, we evaluated the taxonomic status and diversity of local Toxorhynchites species. We collected 147 specimens from artificial and natural containers, yielding 97 cytochrome c oxidase subunit I barcodes from 101 sequenced individuals. A Panama-only Neighbor-Joining analysis revealed four distinct, well-supported clusters (I - IV) exhibiting high mean inter-cluster genetic divergence (4.8 - 20.2%), alongside morphological and ecological segregation. A subsequent global Neighbor-Joining analysis incorporating BOLD and GenBank sequences recovered Toxorhynchites as a strongly supported monophyletic group (99.0% bootstrap support) and revealed deep phylogenetic divergence separating Old World Toxorhynchites (Toxorhynchites) (Clades A and B) from New World Toxorhynchites (Lynchiella) (Clades C - G) lineages. Panamanian Clusters I and II formed two distinct groupings within Clade G, nesting as sisters to Subclade G1 characterized by severe taxonomic discordance involving sequences labeled as Tx. moctezuma, Tx. theobaldi, and Tx. rutilus. Cluster III nested within Tx. hypoptes (1.5% divergence) in Clade E, whereas Cluster IV associated with Tx. haemorrhoidalis s.l. (>5.0% divergence) in Clade C. The high degree of taxonomic uncertainty or cryptic diversity uncovered within Clades C, D, and G underscores an urgent need for formal taxonomic revision to accurately delimit species boundaries. Due to its widespread peri-urban distribution, Cluster I (within the Tx. moctezuma s.l. complex) shows the greatest promise for mass-rearing and Aedes biocontrol applications in Panama.

## Introduction

The genus *Toxorhynchites* (Diptera: Culicidae) is a group of large, diurnal, and colorful mosquitoes primarily found in tropical and subtropical regions around the world (Focks 2007). Unlike most culicid members, female *Toxorhynchites* are non-hematophagous, and therefore harmless to humans in terms of nuisance biting and pathogen transmission (Steffan and Evenhuis 1981). *Toxorhynchites* larvae are voracious predators of other container-breeding mosquito larvae, and as such they have been proposed as potential bio-control agents during integrated vector control campaigns (Steffan and Evenhuis 1981, Collins and Blackwell 2000, Donald et al. 2020). However, the efficacy of practical applications employing *Toxorhynchites* remain hindered by considerable knowledge gaps in their taxonomy, ecology, and geographic distribution (Sukupayo et al. 2024).

*Toxorhynchites* is divided into four subgenera: *Toxorhynchites* (Theobald, 1901), *Afrorhynchus* (Ribeiro, 1992), *Lynchiella* (Lahille, 1904) and *Ankylorhynchus* (Lutz, 1904), which together encompass approximately 91 recognized species globally (Donald et al. 2020). While separating the overarching subgenera is relatively straightforward due to discrete geographic ranges, scientists face significant challenges when trying to identify individual species within them (Steffan and Evenhuis 1981). A prime example of this difficulty occurs within the New World subgenus *Lynchiella*, a group so morphologically identical that adult females can look completely indistinguishable. The cryptic nature of adult *Toxorhynchites*, coupled with the convergent predatory pattern of their larvae, further complicates species identification and classification based solely on morphological characters (Steffan 1975, Collins and Blackwell 2000, Donald et al. 2020). For example, frequent misidentifications and incorrect synonymity of neotropical species such as *Tx. moctezuma sensu lato (s.l.)* (Dyar & Knab, 1906a), *Tx. theobaldi s.l.* (Dyar & Knab, 1906a) and *Tx. hypoptes s.l.* (Knab, 1907) are documented in the scientific literature (Zavortink and Chaverri 2009). Following the work of Dyar (1928) and Theobald (1903), Zavortink and Chaverri (2009) corroborated these three taxa as independent evolutionary units based on morphological characters and historical collection records. However, there is still noticeable disagreement as description of specimens identified as conspecifics, particularly *Tx. moctezuma s.l.*, continue to show conflicting characters (Dyar 1928, Zavortink and Chaverri 2009, Mendez-Andrade et al. 2019, Torres-Avendaño et al. 2021). Furthermore, the practice of lumping specimens into cryptic species complexes or the use of subspecies ranks in their classification (*Tx. haemorrhoidalis* Fabricius, 1787) highlights the need for more rigorous molecular validation approaches (Harbach and Wilkerson 2023).

Refining *Toxorhynchites* taxonomy through molecular phylogenetics is essential to provide a reliable foundation for ecological research. Early studies aiming at establishing the position of this genus within Culicidae utilized partial DNA sequences of the mitochondrial cytochrome c oxidase subunit I gene (COI) (Mitchell et al. 2002). Subsequently, broader phylogenies incorporating additional gene markers and whole mitogenomes offered insights into the monophyletic status of *Toxorhynchites* (Shepard et al. 2006, Reidenbach et al. 2009, Zhou et al. 2014, da Silva et al. 2020, Lorenz et al. 2021). However, these studies limited their scope to include only a few species from a handful of geographic regions. To better understand species limits and evolutionary relationships within *Toxorhynchites*, dedicated studies must incorporate greater taxa diversity from all inhabited regions. Although genetic differentiation has previously helped identify and classify *Toxorhynchites* species on a regional scale (Maquart et al. 2023, Ortega-Morales et al. 2024), such comprehensive molecular validation remains lacking for Mesoamerica.

The “Mosquitoes of Middle America” project reported three *Toxorhynchites* species from the Isthmus of Panama, based on approximately 58 collection records (Heinemann and Belkin 1978). However, because the authors noted that morphological identifications were preliminary, combined with the limited geographic scope of the sampling and a lack of molecular data, this historical work provides only partial insights into local species diversity. In Panama, various molecular markers, including partial sequences of the mitochondrial 3′ COI, 5′ COI, the ribosomal internal transcribed spacer two (ITS2) and the single copy nuclear *white* gene, have been successfully paired with morphology to resolve the taxonomic status of malaria vector mosquitoes (*Anopheles albimanus* Wiedemann, 1820, *Anopheles punctimacula s.l.*, Dyar and Knab, 1906b and *Anopheles triannulatus s.l.* Neiva & Pinto 1922), identifying molecularly distinct lineages across the country (Loaiza et al. 2010, Loaiza et al. 2013, Moreno et al. 2013). To date, similar integrative methodologies have not been applied to delimit *Toxorhynchites* species in Panama or the wider Neotropics, despite their proven utility in refining historical demography and informing vector management strategies.

This study evaluates the taxonomic status of *Toxorhynchites* specimens from Panama. Utilizing a country-wide sampling approach, we integrated adult morphology with COI sequences (Folmer region) to validate species boundaries among previously reported taxa. Furthermore, to provide a broader phylogenetic context, we compared these molecular data against publicly available COI sequences from 10 countries and 3 continental regions archived in the Barcoding of Life Data System (BOLD) (Ratnasingham and Hebert 2007) and GenBank (Benson et al. 2012) databases. Finally, we reviewed geographic distribution patterns and ecological attributes among the Panamanian lineages to gain insights into their viability as biological control agents targeting container-breeding *Aedes* vectors.

## Materials and Methods

### Mosquito Sampling and Species Identification

Sampling was conducted across 10 localities distributed through 8 of the 10 provinces of Panama. We employed oviposition traps (ovitraps) consisting of 16-ounce black plastic containers, 5-liter black plastic buckets, 5-gallon black plastic buckets, and discarded car tires (each containing approximately 2.5 gallons of water). The deployment of dark-colored artificial containers as the surveillance method targeting gravid female mosquitoes is a standard practice and has proven effective for capturing members of the genus *Toxorhynchites* (Yap and Foo 1984, Hoel et al. 2011). Natural water reservoirs (Phytotelmata), including tree-holes, and epiphytic plants (bromeliads and heliconias), were also inspected for *Toxorhynchites* larvae and pupae. Collected larvae were transported to the insectary and reared to the adult stage. Adult *Toxorhynchites* specimens were examined and imaged using a Leica M205 C stereomicroscope equipped with a Leica DMC2900 digital camera and Leica Application Suite (LAS) software package. Specimens were identified to the species level, or the lowest possible taxonomic rank, using morphological keys and original taxa descriptions (Dyar 1928, Zavortink and Chaverri 2009, Méndez-Andrade et al. 2019, Torres-Avendaño et al. 2021). Sampling completeness was evaluated via individual-based rarefaction and extrapolation using the iNEXT package (Hsieh et al. 2016) in R version 4.6.0 (R Core Team 2026). Species abundance metrics were compiled based on counts of molecularly corroborated individuals, and sampling curves were plotted with 95% confidence intervals generated via bootstrap resampling.

### DNA Extraction and PCR Amplification

Total genomic DNA was extracted from individual *Toxorhynchites* specimens using the NucleoSpin Tissue Kit (Macherey-Nagel, Düren, Germany). For each extraction, the right fore- and mid-legs of adults (both sexes) or the fourth abdominal segments of larvae were sliced, placed into sterile 1.5-ml Eppendorf tubes, and homogenized using disposable sterile pestles with 180 µl of cell lysis buffer and 20 µl of Proteinase K. Samples were incubated overnight at 56 °C in a Precision GP15D water bath (Thermo Scientific, Waltham, MA) following the manufacturer’s protocol. DNA quality and concentration were assessed on a NanoDrop Lite spectrophotometer (Thermo Fisher Scientific, Waltham, MA) using 1 µl of the elution buffer as a blank solution and 1 µl of DNA extract from each sample. The resulting DNA extracts were stored at −20 °C.

Prior to PCR amplification, DNA extracts were diluted 1:10 or 1:20 in molecular-grade water depending on the initial concentration (ng/µl). PCR amplification of the Folmer region of the COI gene was performed in a T100 thermal cycler (Bio-Rad Laboratories, Hercules, CA) using the forward primer LCO1490 (5′-GGT CAA CAA ATC ATA AAG ATA TTG G-3′) and reverse primer HCO2198 (5′-TAA ACT TCA GGG TGA CCA AAA AAT CA-3′) (Folmer et al. 1994). PCR reactions were carried out in a 20-µl total volume containing 1× GoTaq Green Master Mix (Promega Corporation, Madison, WI), 0.5 µM of each primer, 0.8 µl of 5% dimethyl sulfoxide (DMSO), 2 µl of diluted DNA template, and nuclease-free water to volume. Thermal cycling conditions consisted of an initial denaturation at 95 °C for 3 min, followed by 35 cycles of denaturation at 95 °C for 30 s, annealing at 51 °C for 1 min, and extension at 72 °C for 1 min, with a final extension phase at 72 °C for 10 min. Positive controls containing *Aedes albopictus* template DNA from central Panama and negative controls containing nuclease-free water were included in every run. Amplification products (∼658 bp) were verified via electrophoresis on a 1.5% agarose gel stained with GelRed Nucleic Acid Gel Stain (Biotium, Fremont, CA) in 1× Tris-borate-EDTA (TBE) buffer at 100 V for 45 min and visualized using a ChemiDoc MP imaging system (Bio-Rad Laboratories). A 100-bp DNA ladder (Promega Corporation) was employed concurrently as a size standard. Successfully amplified products (10 µl) were purified using ExoSAP-IT Express PCR Product Cleanup Reagent (Thermo Fisher Scientific) and submitted to Macrogen, Inc. (Seoul, South Korea) for bidirectional Sanger sequencing (Sanger et al. 1977). The GenBank accession numbers and BOLD codes for our COI gene sequences will be provided upon revision, as the sequences have already been submitted to both databases.

### Molecular Delimitation of *Toxorhynchites* Species from Panama

Partial COI gene sequences obtained from *Toxorhynchites* specimens were edited using Geneious Prime (Kearse et al. 2012) and aligned via the ClustalW algorithm implemented in MEGA12 (Kumar et al. 2024). To identify clusters of COI sequences representing discrete evolutionary units within our Panama-only dataset, we generated a topology using the Neighbor-Joining (NJ) phylogenetic reconstruction method. This approach was selected for its computational efficiency and reliability in clustering large intraspecific datasets characterized by shallow COI divergences (Hernández-Triana et al. 2019). The NJ tree topology was modeled using Kimura 2-parameter (K2P) model, with node support assessed via 1,000 bootstrap replicates. Pairwise genetic distances were calculated in MEGA12 under the K2P model based on a 2% genetic distance threshold to evaluate inter-clusters divergence. For reference and comparative exploration, we included three additional *Toxorhynchites* sequences mined from the Barcoding of Life Data System (BOLD) database: *Tx. moctezuma* (BOLD: MSQQH086-19), *Tx. hypoptes* (BOLD: ASIND3457-12), and *Tx. haemorrhoidalis* (BOLD: MQCHP095-16) (Ratnasingham and Hebert 2007). An *Anopheles albimanus* sequence (BOLD: MOSN10131-23) was used for tree rooting. The geographic distribution of molecularly assigned *Toxorhynchites* specimens from Panama was visualized using R and the ggplot2 package (Wickham 2016), linking each specimen’s cluster assignment to its respective collection locality. The resulting NJ topology served as the initial framework for interpreting *Toxorhynchites* species delimitation within the country.

### Phylogenetic relationships of Toxorhynchites using mined Barcoding Sequences

To place *Toxorhynchites* lineages from Panama within a broader phylogenetic context, we expanded our dataset with mined COI sequences from the BOLD and GenBank databases (Benson et al. 2012) using the tiered selection criteria described in Table 1. This selective approach prioritized high-confidence records to maximize taxonomic diversity and geographic representation within the genus *Toxorhynchites* while minimizing potential misidentification errors. We utilized this expanded dataset to reconstruct an initial phylogeny using the NJ method under the Kimura 2-parameter (K2P) model with 1,000 bootstrap replicates. This initial dataset encompassed our sequences from Panama, 86 mined *Toxorhynchites* sequences, and 5 additional taxa from various other culicid genera. To optimize visualization and reduce sequence redundancy while preserving the topological structure of the initial phylogenetic tree, we generated a second, condensed NJ tree using the same parameters but fewer sequences. This shortened dataset contained representative sequences from each Panamanian lineage and a subset of representative sequences from each mined *Toxorhynchites* lineage. The final global dataset included 12 of our Panamanian *Toxorhynchites* sequences and 37 mined *Toxorhynchites* sequences from Mexico (n = 8), the United States (n = 1), Costa Rica (n = 4), French Guiana (n = 9), Trinidad and Tobago (n = 1), Japan (n = 4), China (n = 6), Thailand (n = 1), Sri Lanka (n = 1), and Australia (n = 2). The resulting multi-regional *Toxorhynchites* phylogeny served as the framework for interpreting global species relationships and confirming natural monophyletic classifications. Pairwise genetic distances for this global dataset were calculated using the same parameters applied to the Panama-only dataset. To evaluate the stability of our global NJ topology, we also performed a complementary Maximum Likelihood (ML) phylogenetic analysis under the K2P substitution model with 1,000 bootstrap replicates.

**Table 1.** The inclusion and exclusion criteria for the selection of COI gene sequences from BOLD and GenBank databases.

| Inclusion Criteria | Exclusion Criteria |
| --- | --- |
| Sequences targeted to the standard Folmer region of the *COI* gene | Non-target genes or *COI* regions outside the Folmer fragment |
| Data retrieved from GenBank or BOLD Systems databases | Sequences obtained from third-party or unverified platforms |
| High-quality sequences without ambiguous bases (Ns) | Low-quality or single-read raw sequences lacking validation |
| Sequence length $\geq 600$ bp | Short fragments ($< 600$ bp) |
| Functional sequences devoid of premature stop codons | Pseudogenes or sequences containing stop codons (NUMTS) |
| Availability of complete accompanying metadata (e.g., country, host) | Incomplete, missing, or corrupted specimen metadata |
| Valid taxonomic identification down to the species level | Sequences restricted to genus, family, or higher taxonomic ranks |

## Results

Initial morphological screening of mosquitoes yielded 147 *Toxorhynchites* specimens collected in 10 sampled localities (Fig. 1). From a processed subset of 101 individuals (92 adults and 9 larvae), we successfully generated 97 high-quality COI sequences, yielding a 96.0% amplification and sequencing success rate. A comprehensive inventory detailed by location code, sample count, morphospecies designations, and corresponding sequencing outcomes is provided in Table 2.

**Fig. 1.**
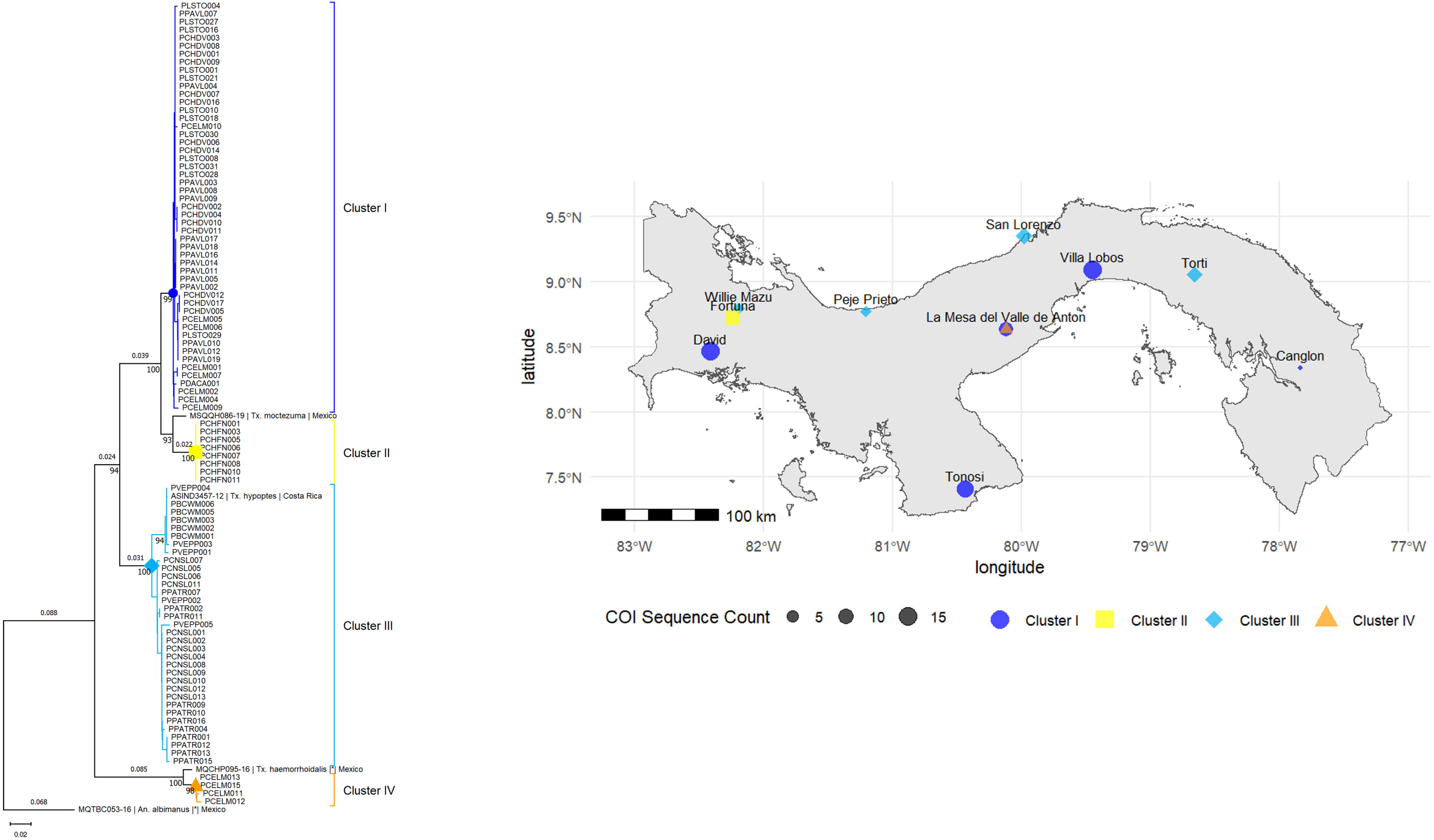
Neighbor-Joining (NJ) phylogenetic tree based on COI gene sequences of *Toxorhynchites* specimens from Panama, alongside a map of sampling localities. Color-coded symbols define the four recovered clusters: Cluster I (blue circle), Cluster II (yellow square), Cluster III (sky-blue diamond), and Cluster IV (orange triangle). Sampling localities on the map correspond to the matching color-coded shapes displayed in the NJ tree.

**Table 2.**
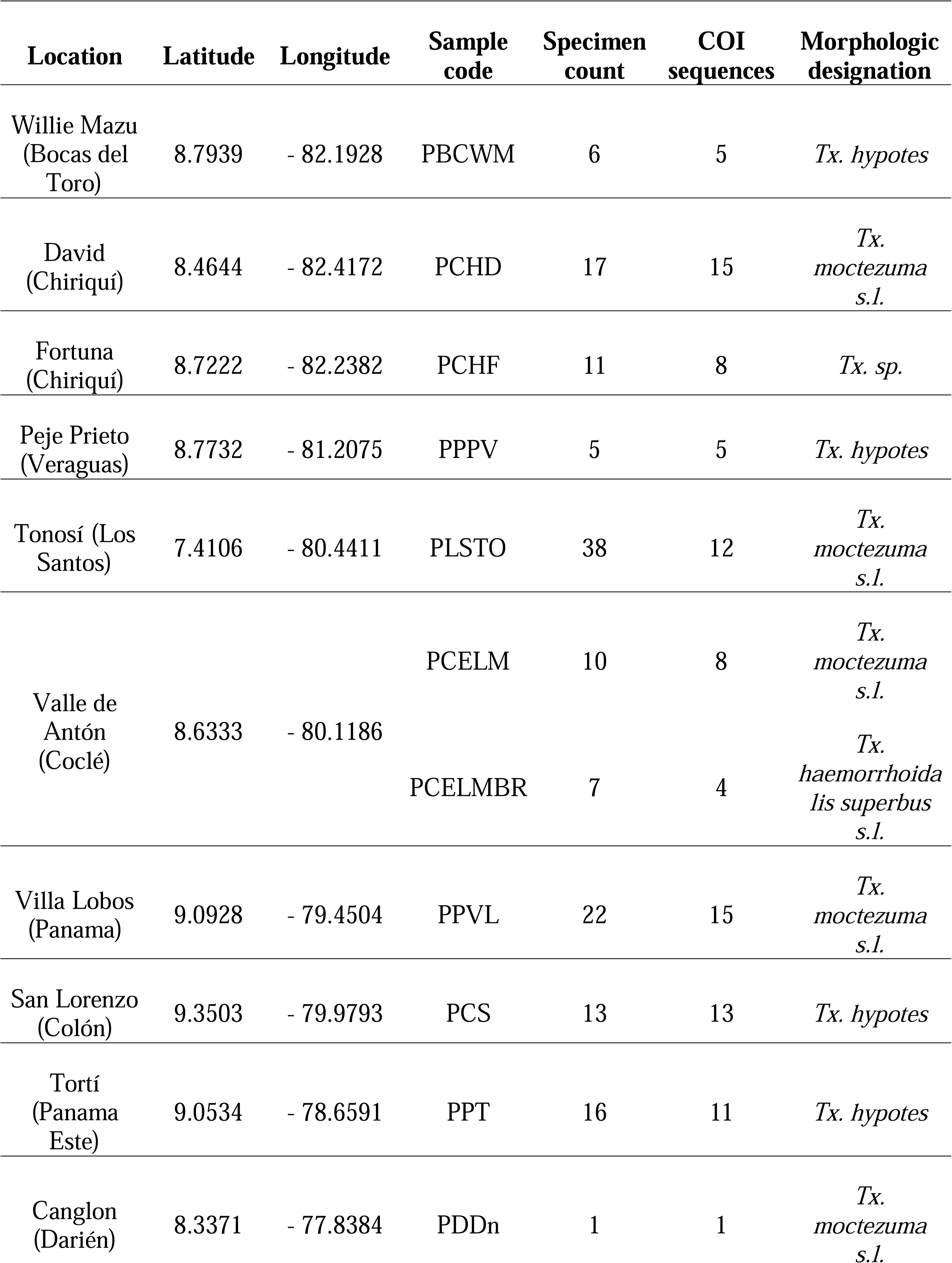
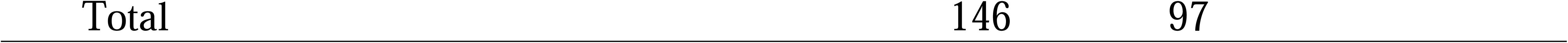
*Toxorhynchites* specimens collected across the country of Panama.

The NJ phylogenetic analysis of Panamanian COI sequences resolved four robustly supported lineages, designated as Clusters I–IV, which correspond exactly to the lineages depicted on the distribution map (Fig. 1). Mean inter-cluster genetic distances ranged from 4.8–20.2%, whereas intra-cluster divergence never exceeded 1.0%. Cluster I comprised 51 COI sequences (25 males, 26 females) and resolved with strong bootstrap support (BS = 99.0%; 0.3% internal divergence); this clade was sister to Cluster II (4.8% divergence) and did not directly group together with any individual reference sequence. Cluster II comprised eight COI sequences (1 male, 1 undetermined, 6 larvae) and resolved with maximal support (BS = 100.0%; 0.0% internal divergence), grouping closely with the *Tx. moctezuma* reference (BOLD: MSQQH086-19) at 3.4% genetic distance. The complete absence of intra-group variation in Cluster II is likely due to a single oviposition event (i.e., a sibling cohort) within the same breeding container. Cluster III comprised 34 COI sequences (20 males, 14 females) and resolved with high nodal support (BS = 100%; 1.0% internal divergence), grouping together with the *Tx. hypoptes* reference (BOLD: ASIND3457-12) at 1.5% genetic distance while remaining distinct from Clusters I and II (9.5% and 11.5% divergence, respectively). Cluster IV comprised four COI sequences (3 males, 1 female) and resolved with high nodal support (BS = 98.0%; 0.3% internal divergence), displaying a 2.1% genetic distance from the *Tx. haemorrhoidalis* reference (BOLD: MQCHP095-16) and representing the most divergent lineage (distances to Cluster I = 18.0%, II = 20.2%, III = 16.0%). Individual-based rarefaction curves asymptoted at the observed sample size (n = 97), and downstream extrapolation yielded no further increase in estimated species richness, indicating that the sampling effort comprehensively captured the available *Toxorhynchites* diversity across the study areas (Supplementary Fig. S1).

*Toxorhynchites* specimens across Clusters I–IV exhibited distinct morphological characteristics that helped distinguish each lineage (Fig. 2). However, poorly preserved specimens introduced significant taxonomic ambiguity, complicating reliable identification. Beyond morphology, these Panamanian clusters displayed noteworthy ecological and microhabitat segregation. Clusters I and III were the most frequently encountered lineages nationwide, demonstrating a sharp contrast in habitat preference; Cluster I occurred in five localities predominantly within lowland peri-urban and anthropogenic environments, whereas Cluster III was restricted to four localities of lowland undisturbed forest habitats. Furthermore, Clusters II and IV were localized to specific high-elevation regions. Cluster II was collected exclusively in Fortuna, Chiriquí, at approximately 1,220 m above sea level (asl), while Cluster IV members were restricted to El Valle de Antón in central Panama at approximately 820 m asl (Fig. 1). This geographic segregation further extended to microhabitat selection: specimens from Clusters I, II, and III were sampled from artificial containers, including ovitraps and discarded tires, whereas Cluster IV individuals were recovered exclusively from bromeliads.

**Fig. 2.**
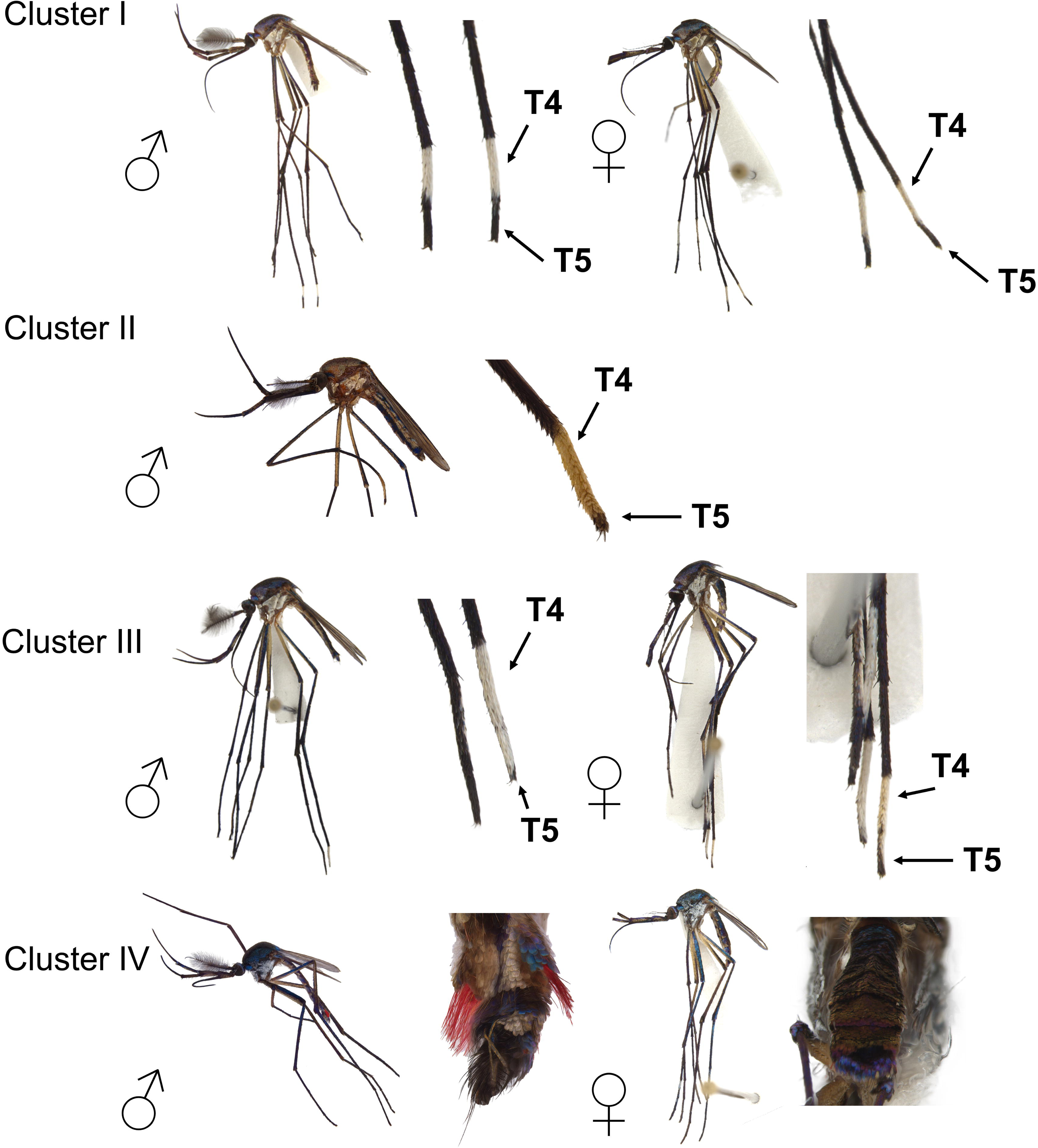
Morphological characteristics of adult *Toxorhynchites* specimens collected in Panama. Cluster I: Male and female hind tarsomeres, showing white fourth tarsomere (T4) and dark fifth tarsomere (T5) in both sexes. Cluster II: Male distal end of hind tarsus, with white scales covering the entire T4 and the base of T5. Cluster III: Male and female hind tarsomeres, with white scales restricted to the exterior surface of T4 and T5. Cluster IV: Male and female distal end of the abdomen.

Out of 273 BOLD COI sequences, only 86 (31.5%) met the inclusion criteria outlined in Table 1 and were utilized for global phylogenetic reconstruction. Of these, 44 sequences also possessed an equivalent accession number in GenBank (Supplementary Table 1). The NJ and ML global phylogenetic reconstructions yielded highly congruent topologies (Supplementary Figs. S2, S3). Consequently, we present only the NJ tree topology for ease of layout, as it preserves all interspecific relationships and nodal support values from the full analysis. The genus *Toxorhynchites* resolved as a robustly supported monophyletic assemblage (99.0% bootstrap support), successfully separated from other culicid genera (Fig. 3). The topology revealed a deep evolutionary divergence (genetic distance 16.8%) separating New World and Old-World lineages, a split that partially corresponds to current subgeneric designation. The Asian and Australasian lineages within the nominate subgenus *Toxorhynchites* (Toxorhynchites) formed two basal clades (A and B). These two broad Australasian and Palearctic groups comprised specimens spanning China, Japan, Thailand, Sri Lanka, and Australia. Clade A grouped *Tx. gravelyi* (Edwards, 1921), *Tx. manicatus yaeyamae* (Bohart, 1956), and *Tx. kempi* (Edwards, 1921) from China and Japan. Clade B split into two distinct subclades: Subclade B1 comprised *Tx. splendens* (Wiedemann, 1819) from China, Thailand, and Sri Lanka alongside *Tx. speciosus* (Skuse, 1889) from Australia, whereas Subclade B2 grouped *Tx. edwardi* (Barraud, 1924) from China and *Tx. towadensis* (Matsumura, 1916) from Japan (Fig. 3). Moreover, American lineages within the subgenus *Toxorhynchites* (Lynchiella) formed a well-supported derived assemblage that diversified into five distinct clades, designated C through G (Fig. 3).

**Fig. 3.**
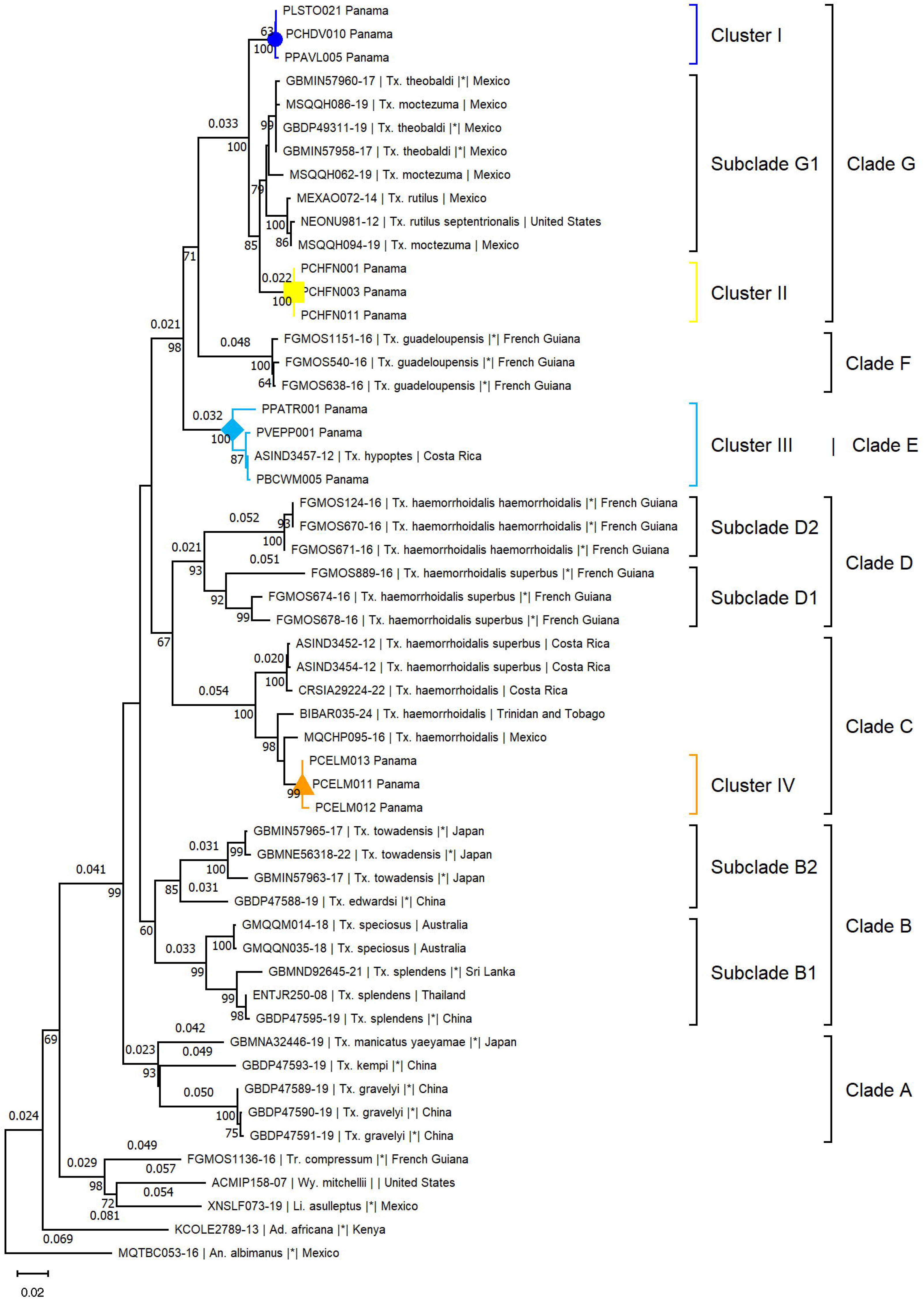
Global Neighbor-Joining (NJ) phylogenetic tree of the genus *Toxorhynchites* based on COI gene sequences, including specimens from Panama and sequences retrieved from BOLD and GenBank. Panamanian lineages are distributed across major clades as follows: Cluster I (blue circle) and Cluster II (yellow square) within Clade G; Cluster III (sky-blue diamond) constituting Clade E; and Cluster IV (orange triangle) within Clade C.

This broad Neotropical and Nearctic group comprised specimens spanning French Guiana, Trinidad and Tobago, Panama, Costa Rica, Mexico, and the United States. Clade C was positioned basally nesting individuals from Panamanian Cluster IV with *Tx. haemorrhoidalis* lineages from Mexico, Costa Rica and Trinidad and Tobago. Clade C resolved as sister to Clade D, which split into two subclades (D1 and D2) containing *Tx. haemorrhoidalis superbus* sequences from French Guiana (Fig. 3). The mean genetic distance separating Clades C and D exceeded 15.4%. Clade E nested Panamanian Cluster III with a single *Tx. hypoptes* reference sequence from Costa Rica. Clade F comprised exclusively sequences of *Tx. guadeloupensis* (Dyar & Knab, 1906) from French Guiana, this lineage remained phylogenetically isolated from all Panamanian Clusters, suggesting it is absent from Panama. Clade G grouped Panamanian Clusters I and II as distinct evolutionary units (exhibiting 4.2% and 3.8% genetic distance to their closest relatives, respectively) sister to another Subclade G1 from Mexico and the United States. Notably, Subclade G1 displayed low internal divergence (mean = 1.6%, maximum = 2.8%) but harbored profound taxonomic discordance, pooling sequences under the names *Tx. moctezuma*, *Tx. theobaldi*, and *Tx. rutilus* (Coquillett, 1896) (Fig. 3).

## Discussion

### Taxonomic Identity of *Toxorhynchites* specimens from Panama

Our local phylogenetic analysis resolved four well-supported clusters defined by robust molecular homology and high bootstrap values, confirming at least four *Toxorhynchites* species in Panama (Fig. 1). Clusters I and II grouped together in the NJ tree but formed two distinct evolutionary units. Cluster II showed the closest genetic affinity to *Tx. moctezuma s.s.* from Mexico, whereas Cluster I formed a separate sister group (Fig. 1). Notably, inter-group divergence between clusters I and II surpass thresholds separating several valid *Toxorhynchites* species in our global phylogeny (Fig. 3). The deep genetic divergence between these two clusters’ mirrors distinct morphological and ecological traits, while highlighting systemic discordances in diagnostic morphological keys. For example, Cluster I specimens displayed diagnostic traits consistent with classical descriptions of *Tx. moctezuma s.s.*, including a white fourth and dark fifth hind tarsomere in both sexes, alongside bluish scutal scales (Dyar 1928, Zavortink and Chaverri 2009). However, they lack the white basal scales on the fifth hind tarsomere diagnostic of *Tx. moctezuma* in some regional descriptions (Méndez-Andrade et al. 2019). Conversely, the single adult from Cluster II that was assessed morphologically, did possess these white basal scales on the fifth hind tarsomere, but exhibited green rather than blue scutal scales. Furthermore, specimens in Cluster I were widely distributed across lowlands areas of Panama, matching historical records for *Tx. moctezuma* near Panama City (Heinemann and Belkin 1978) whereas Cluster II members were restricted to a single high-elevation locality in western Panama near the Costa Rican border, an area with no prior record of *Tx. moctezuma*. The combined morphological, ecological, and molecular evidence strongly suggest that Clusters I and II represent two discrete, undocumented cryptic taxa within the *Tx. moctezuma s.l.* complex rather than simple intraspecific geographic variation within *Tx. moctezuma s.s*.

Panamanian specimens within Cluster III, morphologically identified as *Tx. hypoptes*, nested directly with the species’ sole reference sequence in BOLD (Fig. 1). This specific sequence (BOLD: ASIND3457-12) was exceptionally included despite failing to meet the criteria in Table 1 (i.e., 395 base pairs), ensuring representation for this otherwise unavailable taxon. All Cluster III specimens exhibited diagnostic characters consistent with historical descriptions of *Tx. hypoptes s.s.*, including white scales on the fourth and fifth hind tarsomeres; notably, these scales are restricted to the outer surface in males but occur circumferentially on both surfaces in females (Dyar 1928, Zavortink and Chaverri 2009). Cluster IV specimens were morphologically assigned to *Tx. haemorrhoidalis superbus s.l.*, based on the diagnostic, underdeveloped abdominal patch of red scales (Dyar 1928). This placement was corroborated by their close molecular nesting with a *Tx. haemorrhoidalis* reference sequence from Mexico (Fig. 1). However, our global phylogeny revealed at least five deeply diverging lineages hidden under this single specific taxonomic name (Fig. 3). This polyphyletic pattern strongly supports the systematic remarks of Harbach and Wilkerson (2023), who argued that numerous taxa within the *Tx. haemorrhoidalis s.l.* complex, currently treated as synonyms or subspecies, warrant elevation to independent evolutionary units. Ecologically, the strict association of Cluster IV larvae with bromeliads aligns with historical accounts documenting the *Tx. haemorrhoidalis* complex in Panamanian phytotelmata (Heinemann and Belkin 1978).

### Global Phylogenetic Relationships of *Toxorhynchites*

Our multi-regional phylogenetic analysis provides a robust framework for deciphering the evolutionary landscape of *Toxorhynchites* in Panama and contextualizing it within global lineages. The recovery of *Toxorhynchites* as a monophyletic entity corroborates traditional morphology and recent phylogenomic frameworks, validating the taxonomic distinctiveness of the tribe Toxorhynchitini (Soghigian et al., 2023). At a macro-evolutionary scale, our global topology reveals a deep divergence between Australasian/Palearctic and Neotropical/Nearctic lineages, reflecting historical vicariance and prolonged isolation between Old World *Tx.* (Toxorhynchites) and New World *Tx.* (Lynchiella) groups. Notably, the basal positions and splitting patterns of Clades A and B reveal that the subgenus Toxorhynchites is paraphyletic. In contrast, the nested positions of Clades C–G confirm the monophyly of Lynchiella, suggesting it arose from within a single ancestral Old-World stock.

Regionally, our dataset highlights contrasting systematic challenges across these clades, characterized simultaneously by taxonomic over-splitting and unrecognized cryptic diversity. Old World species within Clades A and B are resolved as cohesive, monophyletic units with negligible intraspecific divergence (< 2.0%) and stable nomenclature. In stark contrast, American Clades C, D, and G display extensive topological inconsistencies and high intraspecific distances, confirming widespread taxonomic misidentifications or unresolved cryptic species complexes (Fig. 3). This taxonomic conflict is manifested in two opposing patterns. First, the low intra-cluster genetic divergence within Subclade G1 suggests that samples identified as *Tx. moctezuma*, *Tx. theobaldi*, and *Tx. rutilus* represent a single, over-split nominal taxon. This pattern aligns with previous reports of extensive diagnostic confusion among these species in Central America (Zavortink and Chaverri 2009). Second, we detected the inverse phenomenon within Clades C and D, including Panamanian Cluster IV; despite being morphologically assigned to the *Tx. haemorrhoidalis* group, these specimens exhibited highly elevated genetic divergence from reference data, signaling hidden, under-split cryptic divergence (Fig. 3).

This high regional complexity is structurally tied to Panama’s geography. The convergence of distinct Panamanian lineages into Clades C, E, and G indicates that the Isthmus operates as a phylogenetic crossroads where northern (*Tx. moctezuma* group) and southern (*Tx. haemorrhoidalis* group) lineages intersect (Fig. 3). This spatial pattern aligns with Panama’s historical role as an Isthmian biological bridge, where recurrent colonization events from isolated Pleistocene refugia shaped modern mosquito community assemblages (Loaiza et al. 2010, Loaiza et al. 2013, Moreno et al. 2013, Loaiza and Miller 2019). However, the resolution of these regional crossroads remains constrained by major global gaps in reference databases. For example, our dataset captured only nine New World species of *Tx. Lynchiella* and seven Old World species of *Tx. Toxorhynchites*, representing minor fractions of the ∼16 and ∼51 described valid species within these respective subgenera (Donald et al. 2020). Crucially, the total absence of sequence data in GenBank or BOLD for the subgenera *Afrorhynchus* and *Ankylorhynchus* prevents an absolute baseline for ancestral node reconstructions. Accordingly, the clades proposed here should be viewed provisionally as phylogenetic species or molecular operational taxonomic units (MOTUs). A greater sequence diversity, particularly from Africa (*Afrorhynchus*) and South America (*Ankylorhynchus*), alongside multi-locus or genomic level markers, will be required to unravel the evolutionary origins and trajectory of the genus *Toxorhynchites*.

### Integrative Taxonomy, Study Limitations, and Future Directions

Our findings underscore that resolving the systematics of *Toxorhynchites* requires a fundamental shift from purely traditional taxonomic paradigms toward a multi-dimensional integrative approach. The diagnostic characters used in species descriptions and morphological keys (Theobald 1903, Dyar 1928) suffer from severe structural constraints because they rely heavily on characters prone to high observer subjectivity and biological variability. Chief among these is the reliance on scale coloration. Because *Toxorhynchites* coloration is a product of physical structural iridescence rather than stable chemical pigmentation, characters like the female fore-tarsi markings shift dynamically with the incident light angle. Consequently, minor viewing adjustments under standard microscopy can easily precipitate erroneous species assignments when applying Dyar’s (1928) criteria. This optical fluidity is further compounded by qualitative, non-quantitative descriptions in the literature, such as "more green than blue" (Dyar 1928) or "pale greenish blue" (Zavortink and Chaverri 2009), which offer no stable diagnostic baseline for modern investigators. Beyond optical limitations, existing morphological keys introduce a severe operational bottleneck due to their reliance on sexually dimorphic and comparative traits. Dyar’s (1928) keys frequently require matched, associated sexes to achieve a confident diagnosis. In modern field surveillance where single, unassociated adults are typically captured, this requirement creates a circular diagnostic failure, rendering isolated specimens unidentifiable without risking extreme observer bias. This issue is critically acute when differentiating sympatric, morphologically ambiguous species complexes (Méndez-Andrade et al. 2019). Similarly, keys relying on comparative rather than discrete, quantifiable traits, such as those separating *Tx. superbus* from *Tx. haemorrhoidalis* based solely on "less developed" caudal setae tufts, are practically unverifiable without immediate access to type-locality morphological vouchers or high-resolution digital photographs. To bypass these interrelated diagnostic hurdles, molecular data must be explicitly linked to comprehensive, single-specimen morphological vouchers (including associated fourth-instar larval and pupal exuviae, paired with the adult features), breeding habitat preferences, and precise geographic location. Importantly, while COI barcodes effectively bridge these morphological gaps, they do not offer a definitive solution. Analytical frameworks must remain resilient against confounding biological and technical artifacts, such as incomplete lineage sorting, nuclear mitochondrial pseudogenes (NUMTs), and public repository misidentifications. This necessity is underscored by the fact that 68.5% of public COI sequences retrieved from BOLD (187 of 273) failed our Table 1 inclusion criteria due to short fragment lengths (<600 bp), premature stop codons, or poor sequence quality. Retaining these anomalous sequences during phylogenetic reconstruction would ultimately compound topological errors.

All these taxonomic ambiguities directly intersect with our sampling limitations and hold major implications for future studies. While localized rarefaction curves indicated sufficient sequencing depth for our sampling effort, our view of Panama’s true *Toxorhynchites* diversity remains constrained by a strong microhabitat collecting bias. Because artificial ovitraps represented our primary sampling mechanism, our dataset inherently favored species adapted to lower-strata, container-like environments, likely missing rare, canopy-dwelling, or high-elevation phytotelm specialists. This sampling gap creates a logical blind spot. Therefore, we cannot fully delineate the evolutionary boundaries or geographic ranges of Clusters I, II, III and IV without sampling the vertically stratified microhabitats, such as canopy bromeliads, epiphytes, and natural tree holes, where these specialized lineages might still reside. Transitioning to a systematic, phytotelm-targeted surveillance protocol will be essential not only to uncover hidden cryptic lineages but also to build the robust, voucher-supported taxonomic baseline required to effectively deploy these predatory mosquitoes as targeted biological control agents against invasive *Aedes* vectors in Panama.

## Conclusions

This study represents the first research effort dedicated to Panamanian mosquitoes to include the genus *Toxorhynchites* since Heinemann and Belkin (1978). Furthermore, it is one of the few in the Americas to implement molecular methods for their taxonomic delimitation. Our results supported a geographically structured evolution for the *Toxorhynchites* genus. Findings further demonstrate that Neotropical *Tx. Lynchiella* diversity remains underestimated when relying solely on morphological criteria. By integrating molecular datasets with classical morphological diagnostics, we identified four distinct species in Panama, uncovered hidden polyphyly within the *Tx. haemorrhoidalis s.l.* complex and revealed potential cryptic species boundaries within *Tx. moctezuma s.l.* complex. The high diversity of *Toxorhynchites* is of particular interest in Panama given the role of larvae as biological control agents of container-breeding *Aedes* vectors. Crucially, Cluster I, identified as a member of *Tx. moctezuma s.l.* complex, was the most abundant, and widely distributed nationwide, exhibiting high adaptation to anthropogenic environments. From an applied perspective, its widespread occurrence and high ecological resilience suggest Cluster I possesses the greatest potential for future colonized mass-rearing and biological control programs in Panama. Future studies incorporating genomic level markers and re-evaluation of voucher morphology (i.e., larvae) will be essential to resolve the taxonomic uncertainties identified in Panamanian lineages and to confirm whether current subgeneric boundaries truly represent natural evolutionary groups.

## Acknowledgments

Special thanks are due to STRI and INDICASAT AIP for their administrative backing, technical guidance, and logistical oversight. We are highly grateful to José R. Rovira and Ronald Peña (INDICASAT AIP) for their expert support in mosquito sampling, rearing, and taxonomy. We also acknowledge the Panamanian Ministry of Environment (MiAmbiente) for authorizing and supporting our scientific fieldwork.

## Funding

Economic support was provided by the National Secretariat of Science, Technology, and Innovation (SENACYT – APY-NI-2023A-08 – New Researchers and Entrepreneurs 2024) to RB. J.R.L.’s research activities are supported by the National System of Investigation of SENACYT (SNI 056–2023 and 002–2026). The funders had no role in study design, data collection and analysis, the decision to publish, or the preparation of the manuscript.

---

**Supplementary Table S1.**
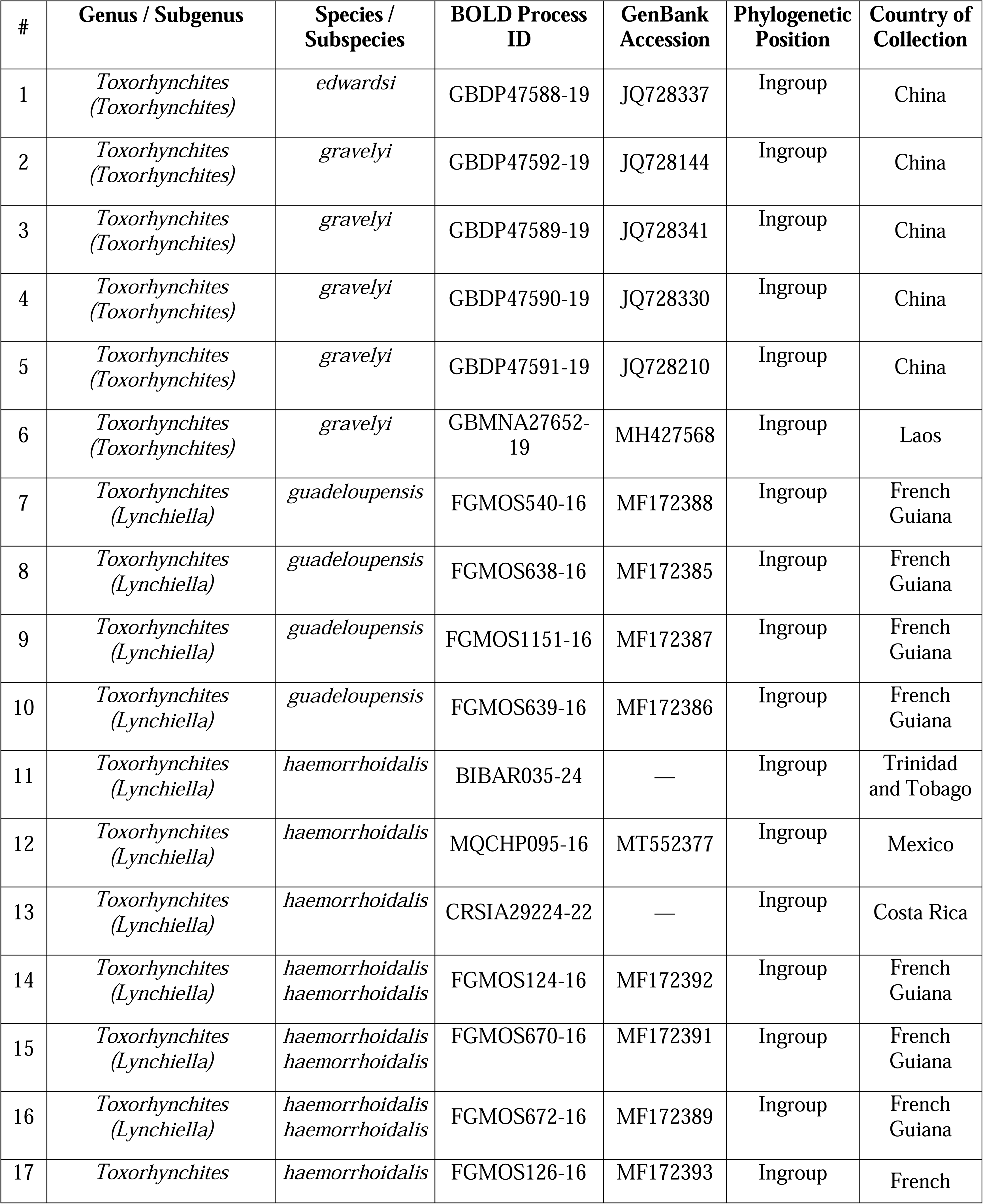
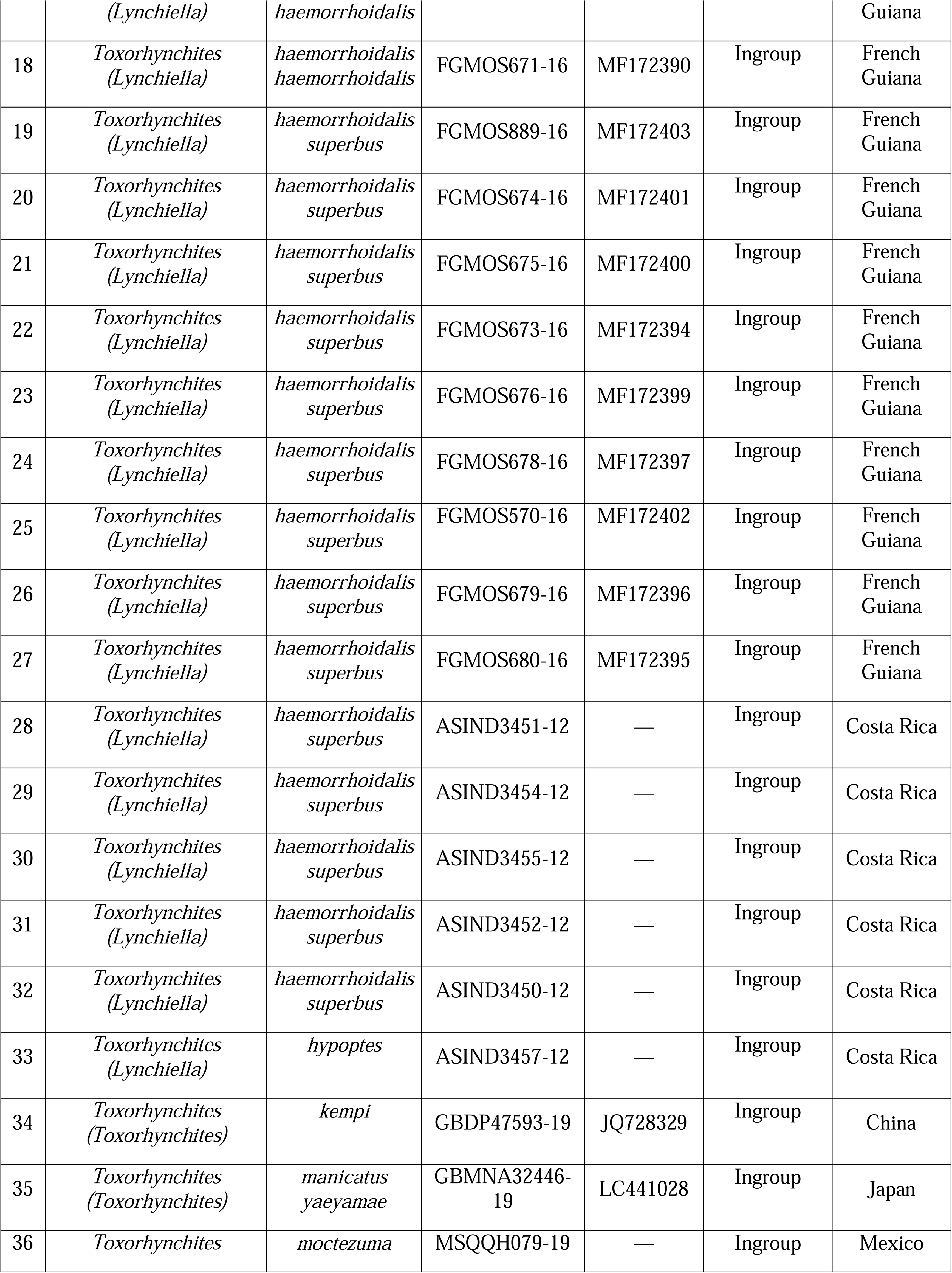
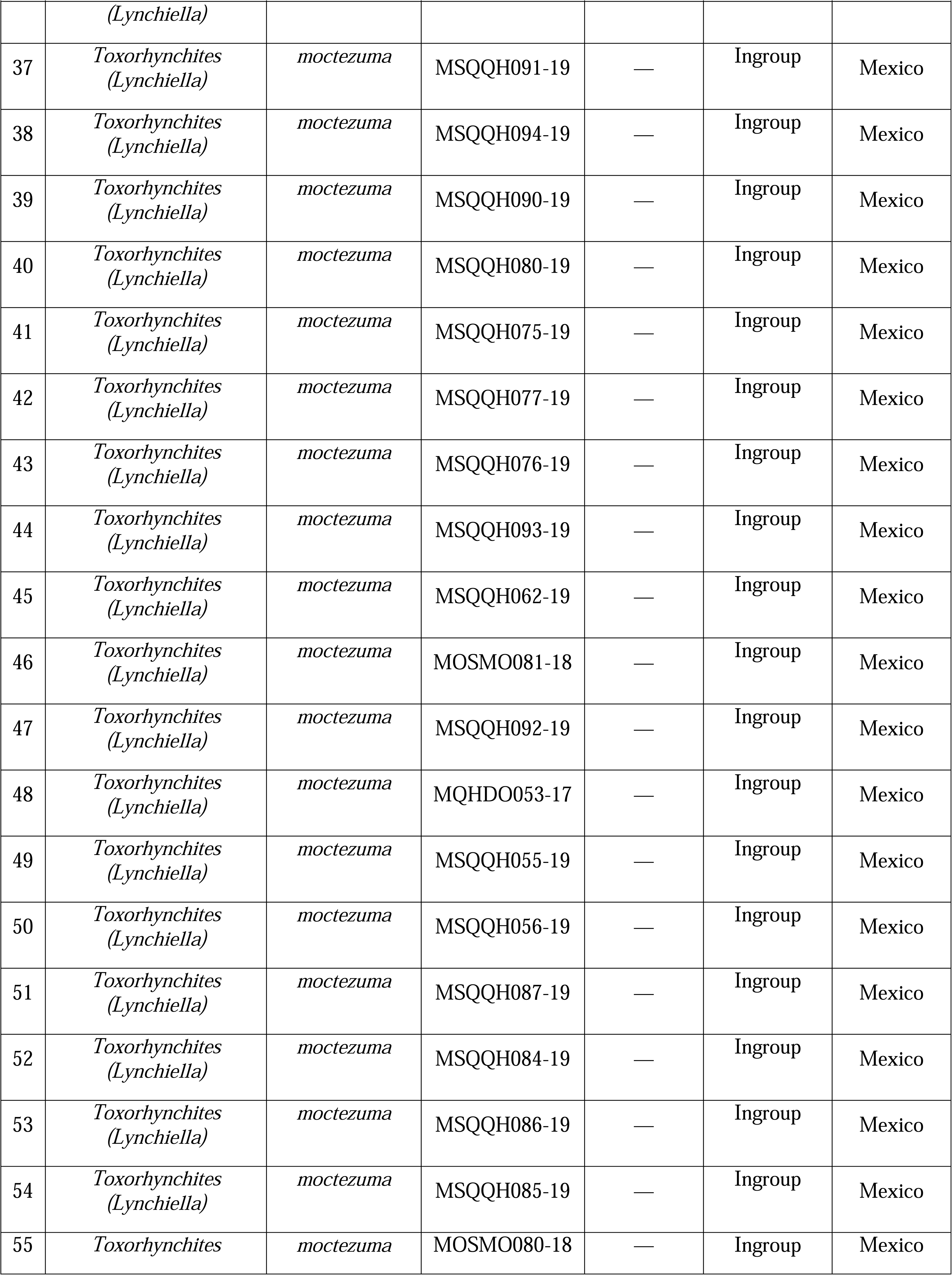
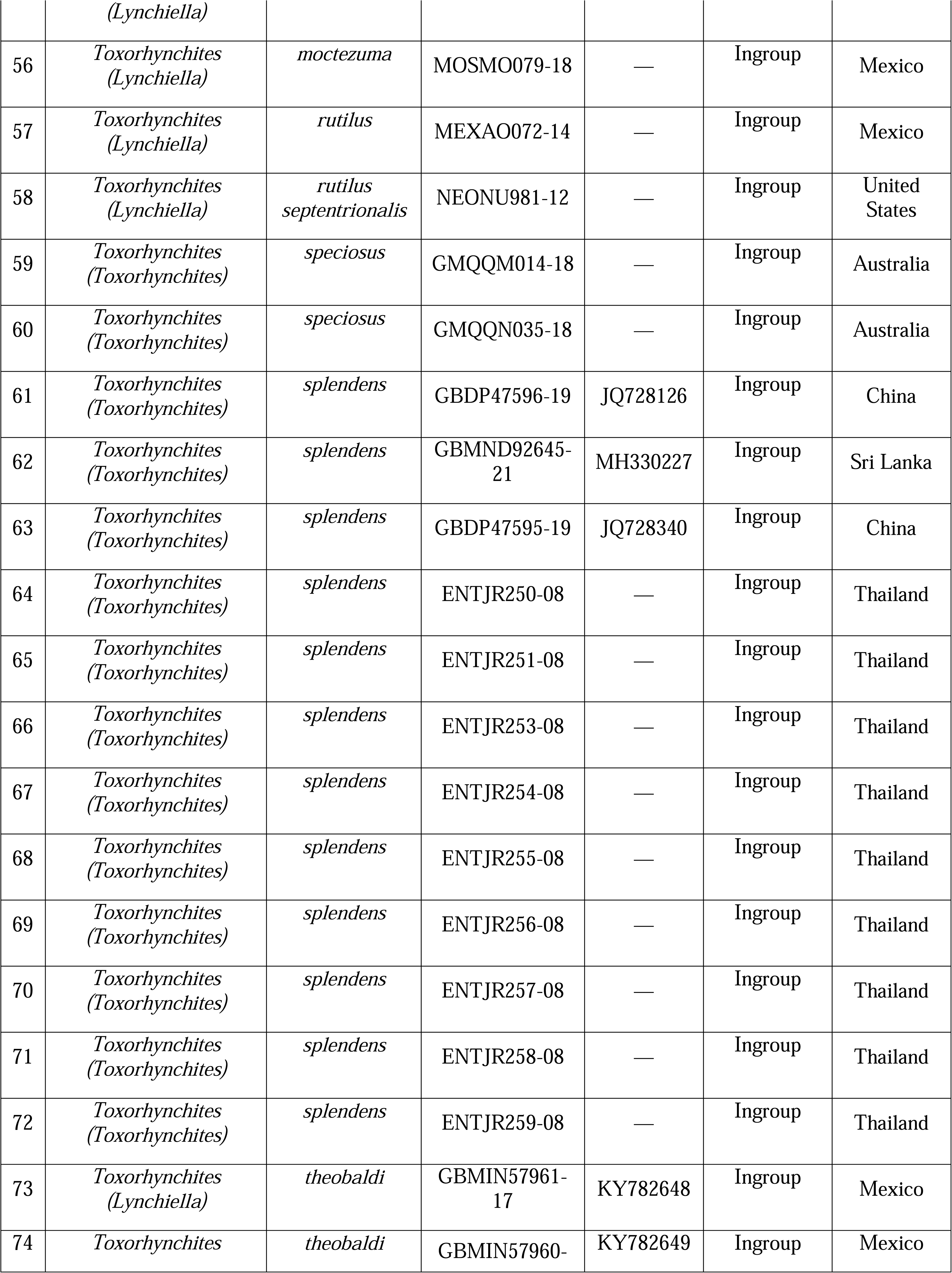
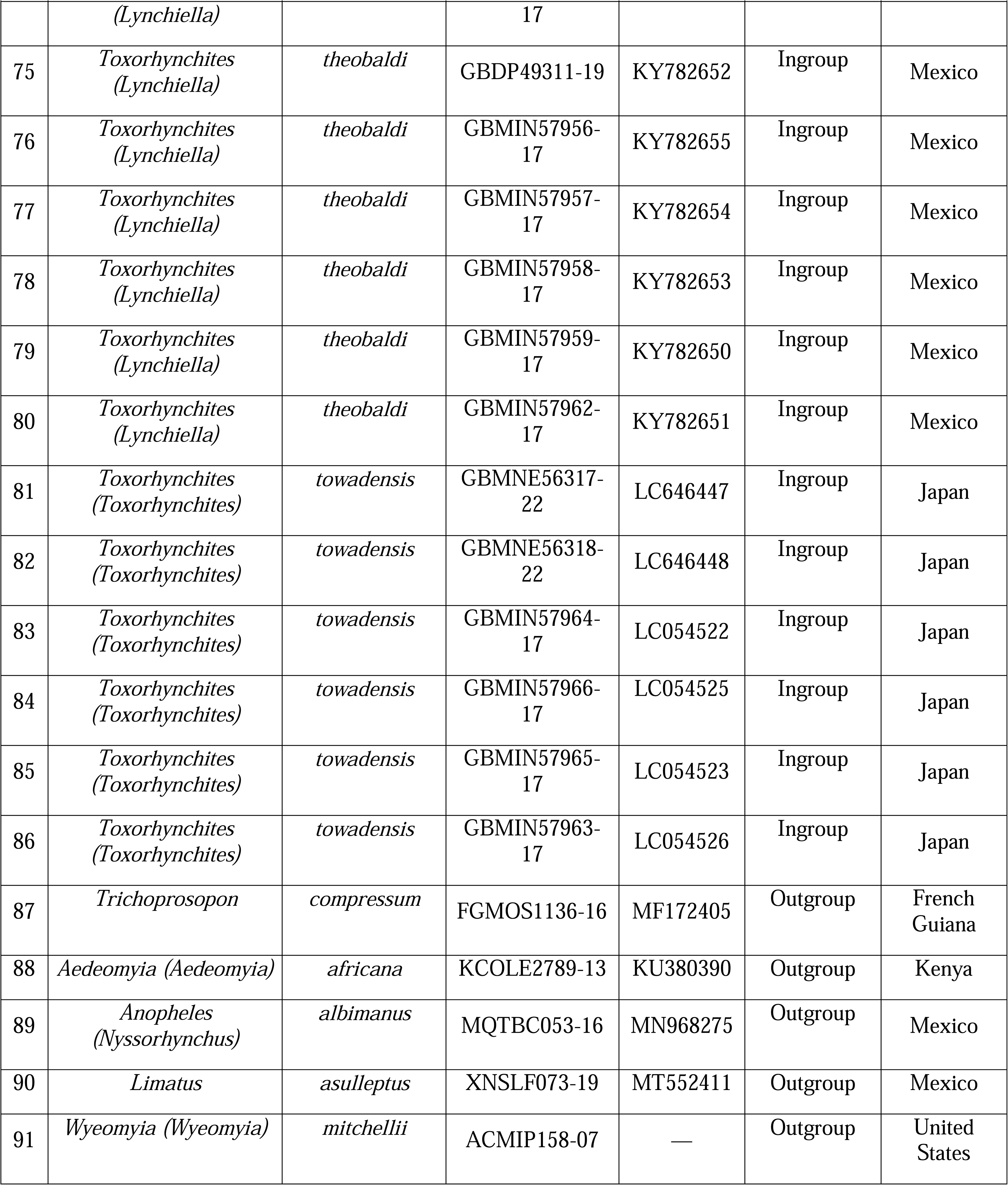
Taxon nomenclature, BOLD process IDs, GenBank accessions, phylogenetic position, and collection countries for the mined COI sequences included in the global *Toxorhynchites* phylogenetic analysis. Missing GenBank accessions are indicated by an em dash (—).

**Supplementary Figure S1.**
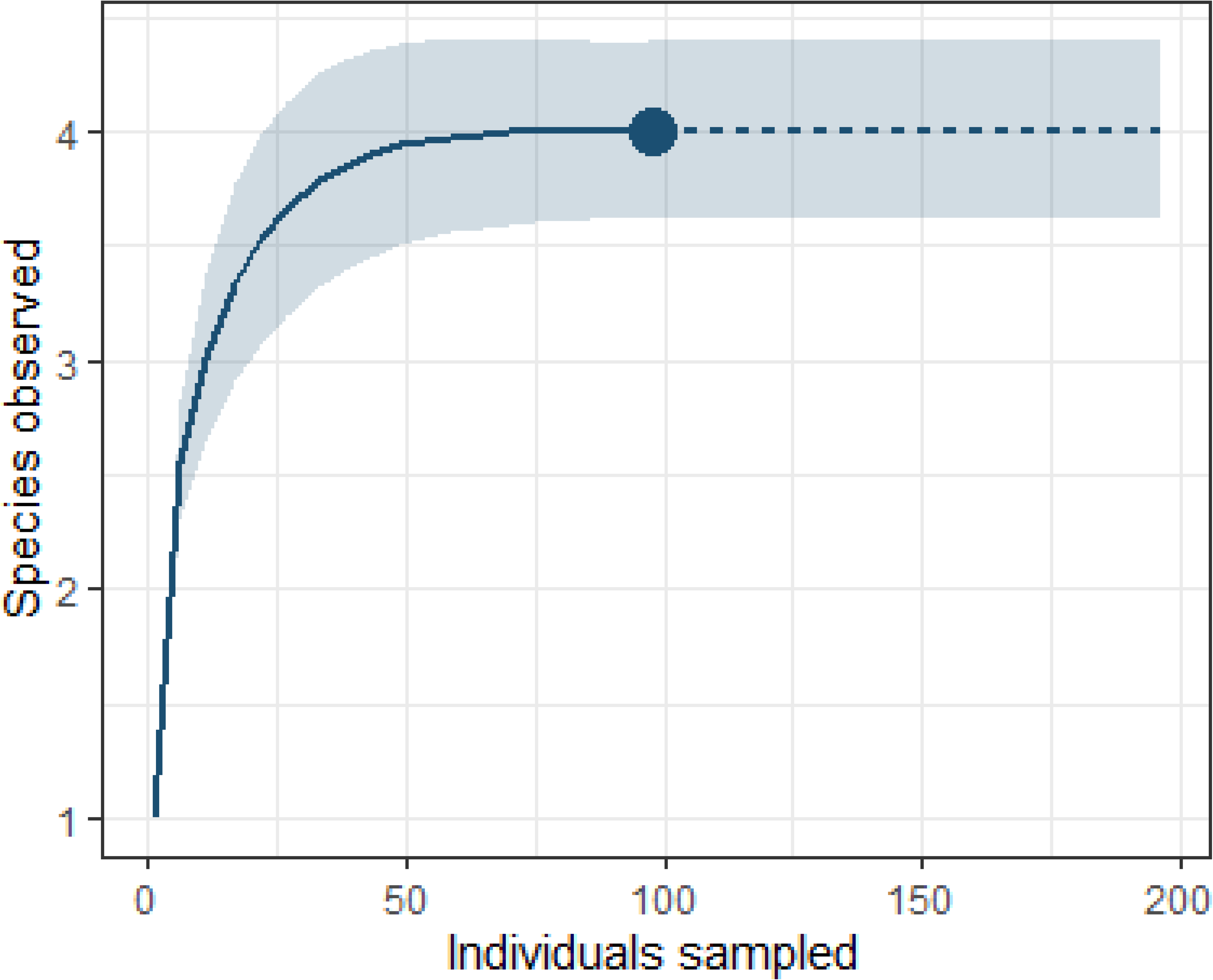
Individual based rarefaction and extrapolation curve for the *Toxorhynchites* assemblage based on the molecularly confirmed individuals. The solid line represents the rarefaction estimate, the dashed line shows the extrapolation, and the shaded region indicates the 95% confidence interval. The filled point marks the observed richness at 97 specimens.

**Supplementary Figure S2.**
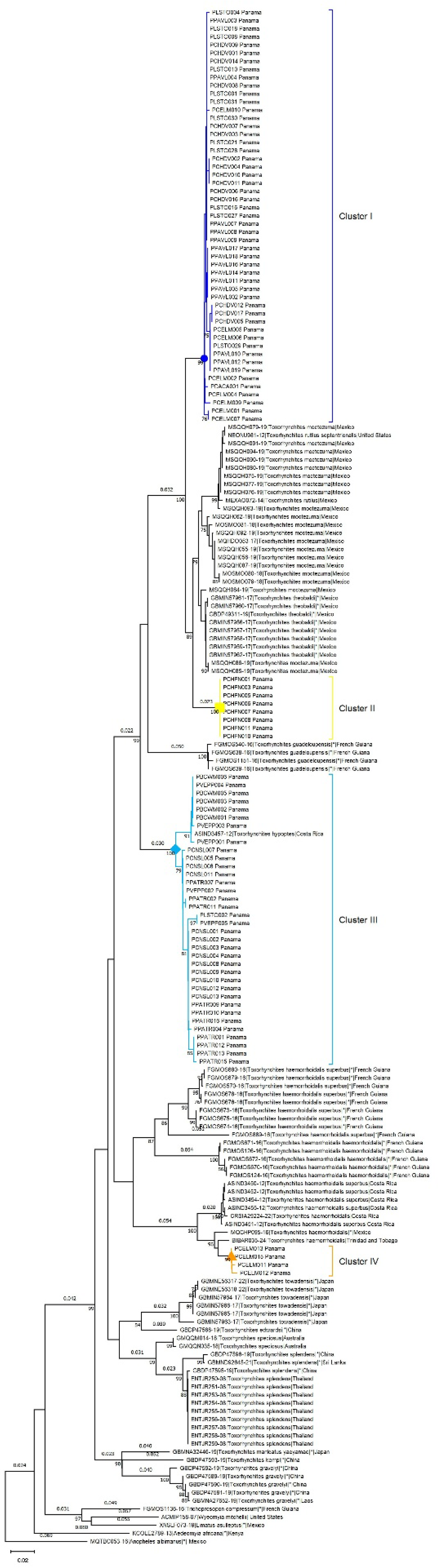
Global Neighbor-Joining (NJ) phylogenetic tree based on COI gene sequences, including 97 *Toxorhynchites* specimens from Panama, 86 reference sequences retrieved from public databases, and 5 outgroup taxa from other culicid genera. Panamanian lineages are indicated by color-coded symbols: Cluster I (blue circle), Cluster II (yellow square), Cluster III (sky-blue diamond), and Cluster IV (orange triangle).

**Supplementary Figure S3.**
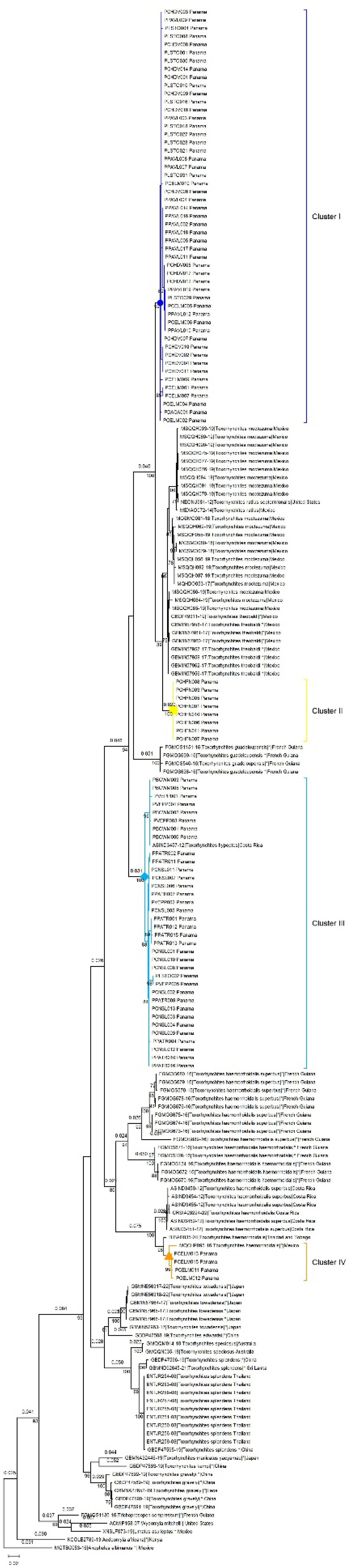
Global Maximum-Likelihood (ML) phylogenetic tree based on COI gene sequences, including 97 *Toxorhynchites* specimens from Panama, 86 reference sequences retrieved from public databases, and 5 outgroup taxa from other culicid genera. Panamanian lineages are indicated by color-coded symbols: Cluster I (blue circle), Cluster II (yellow square), Cluster III (sky-blue diamond), and Cluster IV (orange triangle).

## Notes

### Competing Interest Statement

The authors have declared no competing interest.

